# A Software Toolkit for TMS Electric-Field Modeling with Boundary Element Fast Multipole Method: An Efficient MATLAB Implementation

**DOI:** 10.1101/2020.02.09.941021

**Authors:** Sergey N. Makarov, William A. Wartman, Mohammad Daneshzand, Kyoko Fujimoto, Tommi Raij, Aapo Nummenmaa

## Abstract

**Background:** Transcranial magnetic stimulation (TMS) is currently the only non-invasive neurostimulation modality that enables painless and safe supra-threshold stimulation by employing electromagnetic induction to efficiently penetrate the skull. Accurate, fast, and high resolution modeling of the electric fields (E-fields) may significantly improve individualized targeting and dosing of TMS and therefore enhance the efficiency of existing clinical protocols as well as help establish new application domains.

**Objective:** To present and disseminate our TMS modeling software toolkit, including several new algorithmic developments, and to apply this software to realistic TMS modeling scenarios given a high-resolution model of the human head including cortical geometry and an accurate coil model.

**Method:** The recently developed charge-based boundary element fast multipole method (BEM-FMM) is employed as an alternative to the 1st order finite element method (FEM) most commonly used today. The BEM-FMM approach provides high accuracy and unconstrained field resolution close to and across cortical interfaces. Here, the previously proposed BEM-FMM algorithm has been improved in several novel ways.

**Results and Conclusions:** The improvements resulted in a threefold increase in computational speed while maintaining the same solution accuracy. The computational code based on the MATLAB^®^ platform is made available to all interested researchers, along with a coil model repository and examples to create custom coils, head model repository, and supporting documentation. The presented software toolkit may be useful for post-hoc analyses of navigated TMS data using high-resolution subject-specific head models as well as accurate and fast modeling for the purposes of TMS coil/hardware development.

## 1. Introduction

Neuropsychiatric disorders are a leading source of disability and require novel treatments that specifically target the mechanisms of disease. As such disorders are thought to result from aberrant neuronal circuit activity, neuromodulation approaches are of increasing interest given their potential for manipulating circuits directly (Bikson et al 2018). Noninvasive, noncontact transcranial magnetic stimulation (TMS), which uses magnetic induction to generate current internal to the brain remotely via a coil placed next to the subject’s head, is one of the currently used major neurostimulation modalities (Kobayashi and Pascual-Leone 2003, Rossi et al 2009, McMullen 2017). More recently, the same physical principles have been also employed in embedded microcoils targeting selected populations of neurons while avoiding problems associated with the tissue-electrode interface (Bonmassar et al 2012, Lee et al 2016, Lee and Fried 2016). Due to the non-invasive nature of TMS, computational modeling of the electric fields within a patient-specific head model is the major and often only way to foster spatial targeting and/or obtain a quantitative measure of the stimulation intensity.

While several alternatives exist (Sim4Life, ANSYS Maxwell), the predominant FEM-based TMS modeling software is currently SimNIBS v. 1-3 (Thielscher et al 2015, Opitz et al 2015, Nielsen et al 2018, Saturnino et al 2019a, Saturnino et al 2019b, Saturnino et al 2019c). This software uses robust formulations of the finite element method (FEM). In switching from the open-source 1st order FEM solver getDP to a more rigorous 1st order FEM formulation enabled by SimNIBS 3.0, the software achieves a remarkable performance improvement: an iterative FEM solution computed in less than 30 sec using a head model with a nodal density of 0.5 nodes/mm^2^, processed on an Intel i7-7500U laptop processor (2 cores) with a clock speed of 2.7-3.5 GHz (Saturnino et al 2019b, Saturnino et al 2019c).

In this article, we present an alternative modeling approach for fast, high-resolution modeling of transcranial magnetic stimulation (TMS). The mathematical algorithm is based on the direct formulation of the boundary element method in terms of induced charge density at the interfaces naturally coupled with the fast multipole method or BEM-FMM originally described in (Makarov et al 2018, Htet et al 2019a). Some distinct features of the BEM-FMM based modeling approach developed herein include:

i. High numerical accuracy, which was recently shown to exceed that of the comparable finite element method of the first order (Gomez et al 2019).
ii. Unconstrained field resolution close to and across cortical surfaces, including both the outer cortical surface (the interface between gray matter (GM) and cerebrospinal fluid (CSF) following terminology of Li et al 2012 and the inner cortical surface (the interface between white matter (WM) and gray matter (GM)). Since the solution is fully determined by the conductivity boundaries, the BEM-FMM field resolution within the cortex is not limited by the FEM volumetric mesh size and may reach a micron scale if desired.
iii. Zero post-processing time for the normal components of the electric field close to and across cortical interfaces, once the solution for the induced surface charge density is known.
iv. Comparable speed. For a head segmentation with approximately 1 M facets (default example *Ernie* of SimNIBS 3.x), the improved BEM-FMM algorithm computes the complete numerical solution in approximately 38 seconds (excluding preprocessing time which occurs once per model), while the SimNIBS takes 32 seconds for the matrix solution step alone on the same 2.1 GHz multicore server.
v. Scalability to large-scale / high-resolution models. A surface model with 70 M facets has been considered and computed within two hours, demonstrating the vast potential that the method has to solve large-scale and/or high-resolution problems.
vi. Precise coil modeling and optimization. By employing the fast multipole method, it is possible to model and optimize off-the-shelf and/or custom-designed coil CAD models composed of hundreds of thousands of elementary current elements (Makarov et al 2019).

The first goal of this article is to provide a detailed description of the BEM-FMM numerical algorithm, including several critical improvements, in the Materials and Methods section. The section also details the procedure for importing a head model, the head models available with the software, built-in surface remeshing tools, and NIfTI viewer tools.

The second goal of the study is to demonstrate the resulting method’s speed, accuracy, and resolution, and illustrate the method’s capabilities based on several realistic TMS scenarios given a high-resolution head model including gyral/sulcal folding patterns and a precise coil model (the Results section). Particular attention is paid to electric fields in the vicinity of the inner cortical surface (the white-gray matter interface). The normal field just inside the inner cortical surface (which is significantly higher than the field just outside) and the normal field discontinuity (whose meaning is discussed later) may stimulate either straight or bent pyramidal axons of the fast-conducting pyramidal tract neurons, resulting in D (direct) wave generation (Salvador et al 2011, Lazzaro and Ziemann 2013).

Finally, Appendix A describes the developed software package and walks a potential user through specific computation steps pertinent to one of the study examples. The complete computational code, along with supporting documentation, is available for academic purposes via a Dropbox repository (Dropbox 2019).

## 2. Materials and Methods

In the past, the BEM-FMM algorithm was successfully applied to the modeling of high-frequency electromagnetic (see Song et al 1997, Chew et al 2001) and acoustic (see Chen et al 2008, Burgschweiger et al 2012a, Burgschweiger et al 2012b, Burgschweiger et al 2013, Wu et al 2013) scattering and radiation problems in non-medical fields with a focus on defense applications. The successful implementation of the method for quasistatic bioelectromagnetic problems, however, was lacking. One such implementation was suggested in Makarov et al 2018, Htet et al 2019a, based on accurate coupling of the canonic general-purpose fast multipole method (Greengard and Rokhlin 1987, FMM 2017, Gimbutas et al 2019) and the direct (without using reciprocity) quasistatic boundary element method formulated in terms of induced surface charge density, also known as the adjoint double layer formulation (Barnard et al 1967, Rahmouni et al 2018). Below, we describe the complete BEM-FMM algorithm along with its most recent improvements and establish the method’s convergence.

### 2.1. Direct charge-based boundary element method in a conducting medium

Induced charges with a surface charge density *ρ*(***r***) in C/m^2^ reside on macroscopic or microscopic tissue conductivity interface(s) *S* once an external electromagnetic stimulus (a primary electric field ***E**^p^*(***r***), either conservative or solenoidal) is applied. The induced surface charges alter (typically block and/or redirect) the primary stimulus field. The total electric field anywhere in space except the charged interfaces themselves is governed by Coulomb’s law

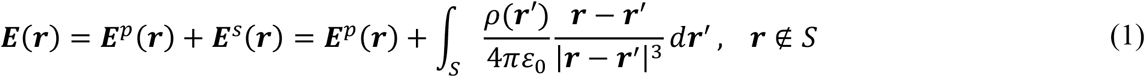

where ε_0_ is dielectric permittivity of vacuum. The electric field is discontinuous at the interfaces. When approaching a charged interface *S* with a normal vector ***n*** from either direction (inside or outside with regard to the direction of the normal vector), the electric field is given by

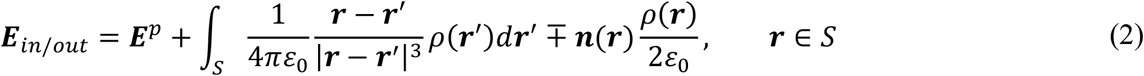

An integral equation for *ρ*(***r***), which is the Fredholm equation of the second kind, is obtained after substitution of Eq. (2) into the quasistatic boundary condition, which enforces the continuity of the normal current component across the interface, that is

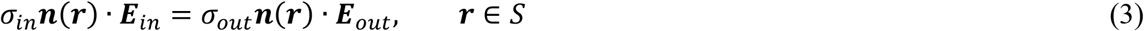

The result has the form

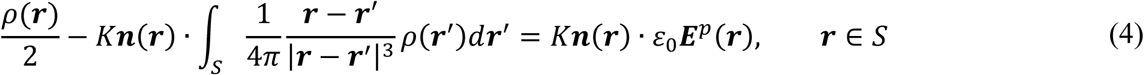

where the electric conductivity contrast 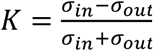 is defined at the interface(s). Here, σ*_in_*, σ_out_ are the conductivities inside and outside with regard to the direction of the normal vector, respectively.

### 2.2. Effect of dielectric permittivity

If we solve Eq. (4) and then substitute the result for *ρ*(***r***) in Eq. (1), the normalization constant σ_0_ will cancel out. Therefore, its exact value does not matter for the subsequent analysis. However, if the displacement currents are significant, extra bound polarization charges will reside on the interfaces (Makarov et al 2015). Their effect is taken into account by considering a complex conductivity in the form σ→ σ+ *j*ωε for a harmonic excitation with angular frequency ω in Eq. (4).

### 2.3. Treatment of interfaces

If the surface is a 2-manifold object with no contact to other surfaces (a “nested” topology where each of the surfaces is associated with a single unique exterior compartment), ***n*** is simply the outer normal vector to the surface; σ*_in_* is the conductivity inside the object; and σ*_out_* is the conductivity of the surrounding medium. If two objects (1 and 2) are in contact with each other as shown in Fig. 1, the joint interface between them should be counted *only once*. In Fig. 1, this interface is counted only for object 1 with *σ*_1_ being the inner conductivity and σ_2_ being the outer conductivity (in the direction of the normal vector ***n***_1_). Facets of object 2 at the interface are now ignored to avoid double-counting. Alternatively, the interface may belong only to object 2, with the direction of the normal vector and the conductivity values switched. In that case, facets of object 1 at the interface would be ignored.

**Fig. 1.**
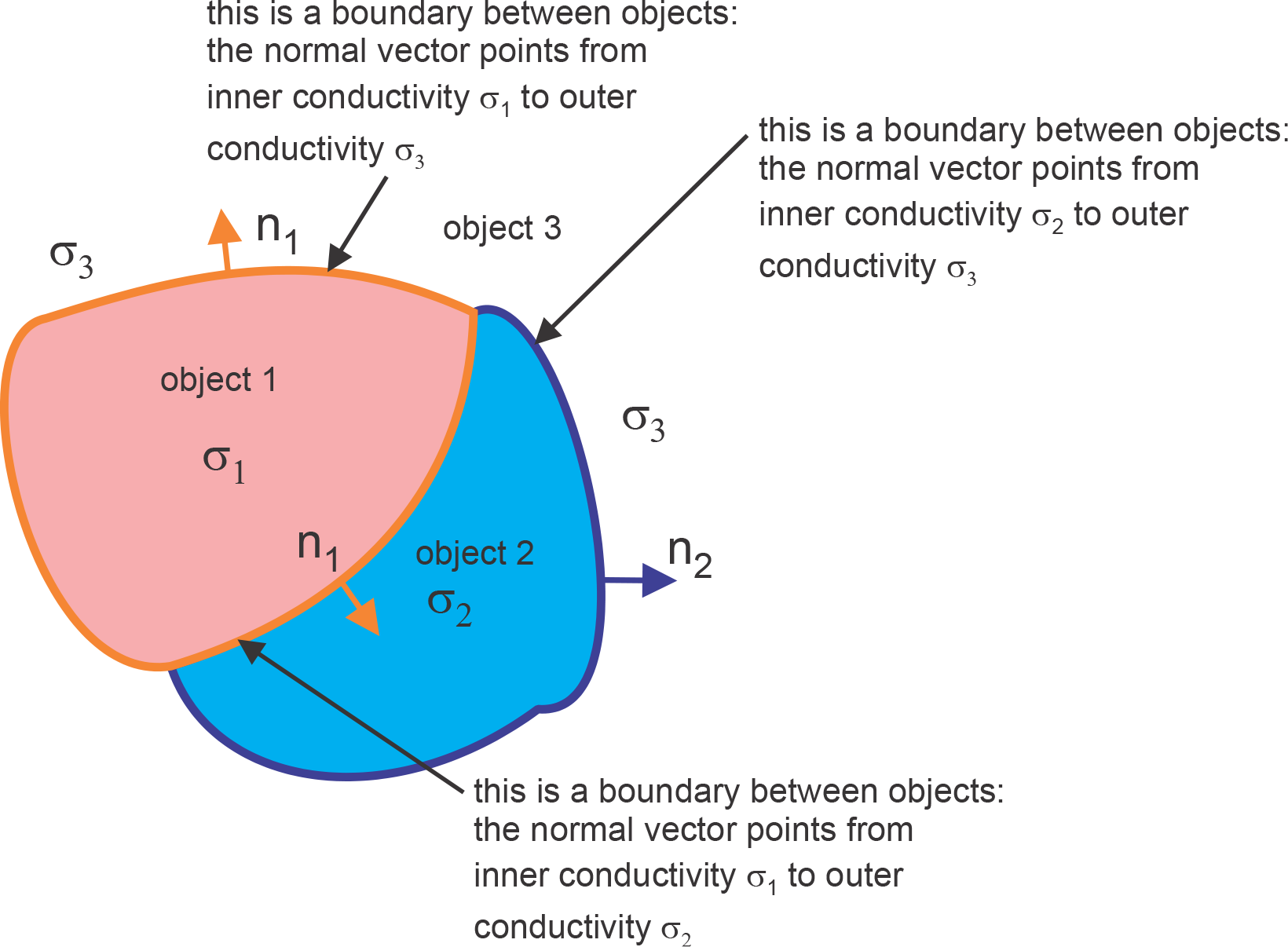
For two objects (1 and 2) in contact with each other, the joint interface between them should be counted *only once*. In the figure, this interface is counted only for object 1 with *σ*_1_ being the inner conductivity and σ_2_ being the outer conductivity (in the direction of the normal vector ***n***_1_). Facets of object 2 at the interface are ignored to avoid double-counting.

From the formal point of view, one needs a composite mesh *without* double coincident facets. Then, for an arbitrary triangular facet *t_m_* of the mesh with a given unit normal vector ***n**_m_*, one needs to know the conductivity σ*_m,out_* “outside” (i.e. in the direction of ***n**_m_*) and the conductivity σ*_m,in_* “inside” (i.e. in the opposite direction of ***n**_m_*). This information is sufficient to completely describe the model.

### 2.4. Normal electric fields at the interfaces

A significant and previously unnoticed advantage of the above approach is an ability to precisely obtain electric fields normal to the cortical surfaces (or any other interfaces) without additional computational cost or postprocessing. Only the solution for the surface charge density is necessary. After taking the scalar product of Eq. (2) with the surface normal vector ***n***, Eq. (4) may then be substituted to explicitly find the normal electric field just inside the surface, ***n***⋅ ***E****_in_*(***r***); the normal electric field just outside the surface, ***n***⋅ ***E****_out_*(***r***); and the normal field discontinuity for any conducting interface, *d**n***⋅ ***E***. All three quantities are directly proportional to each other. One has, for any observation point **r** ∈ *S*,

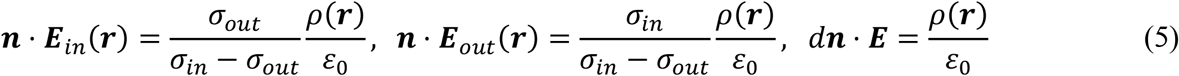

for any conducting interface *S*. We should note that the normal component of the total E-field just inside/outside conductivity boundaries depends explicitly only on the induced surface charge density.

### 2.5. Charge conservation law

The charge conservation law is not explicitly included in Eq. (4); it must be enforced in the form

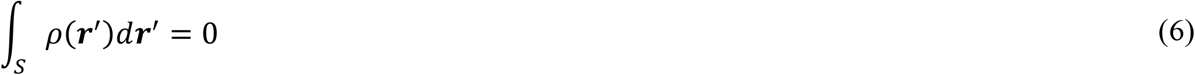

In Eq. (6), *S* is now the combination of all interfaces. Proper implementation of Eq. (6) implies a direct combination with Eq. (4) with a proper weighting; it provides a significantly better and unconstrained convergence rate of the iterative solution, as shown in Appendix A; it also prevents excess charge accumulation at sharp corners of the model.

### 2.6. Model discretization

The surface charge density is expanded into pulse bases (zeroth-order basis functions) on triangular facets *t_m_* with area *A_m_*. The charge density is thus constant for every facet. The Petrov-Galerkin method is then applied to Eq. (4) which gives us a system of *M* linear equations for unknown expansion coefficients *c_m_* in the form

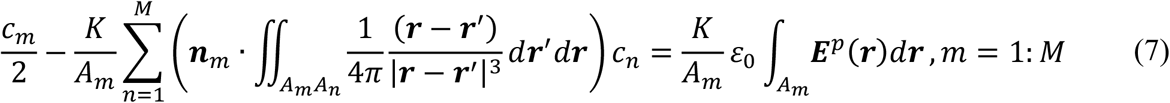

The double potential integrals present in Eq. (7) require care in their numerical evaluation. Facets which are spatially close to one another (i.e., not considered well separated on the lowest level of the FMM octree) cannot be treated with the FMM. These nearfield potential integrals are instead directly calculated and stored in the sparse nearfield BEM matrix using analytical integration for the inner integral and a Gaussian quadrature of 10^th^ degree of accuracy for the outer integrals (Htet et al 2019a). The number of geometrical (based on Euclidian distance) neighbors in Eq. (7) may vary, but a relatively small number may be adequate. It must be noted that these geometrical neighbors may belong to different tissue compartments.

### 2.7. Fast multipole method

The general-purpose fast multipole method (FMM) and its most recent freeware distribution (Gimbutas et al. 2019) is applied to compute the remainder of the integrals of type defined by Eq. (7) using the center-point approximation at face centers ***r****_m_*, yielding

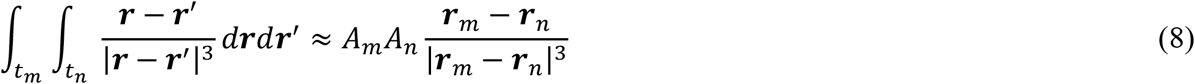

This problem is equivalent to finding the electric field at target points ***r****_m_* generated by the point charges located at source points ***r****_n_*. The accuracy of the FMM (the number of levels) is conventionally estimated for arbitrary volumetric charge distributions. However, for surface-based charge distributions, a much better relative accuracy is observed. For example, with the intrinsic method accuracy set as *ε*= 0.1, the mean error for the pial cortical surface (gray matter shell) may be as low as 0.1% with respect to the electric field amplitude and 0.08 deg with respect to the field angle deviation as compared to the most accurate solution (i.e., the solution where FMM precision is set to maximum).

### 2.8. Primary TMS field

An arbitrary TMS coil is modeled in the form of a very large number of small straight elements of current *i*_j_ (*t*) with orientation vector ***s****_j_* and center coordinate vector ***p****_j_*. Those elements can be either uniformly (Litz wire) or non-uniformly (skin effect) distributed over every conductor’s cross-section. The magnetic vector potential ***A****^p^* of a current element with orientation ***s****_j_* and position ***p****_j_* at an observation point ***C****_i_* is given by (Balanis 2012)

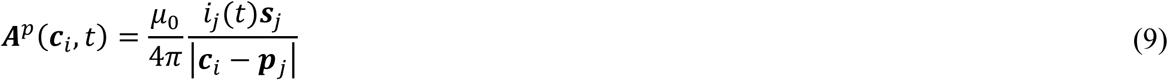

where *μ*_0_ is magnetic permeability of vacuum and index *p* denotes the primary field. The corresponding solenoidal electric field is ***E**^p^* = − *∂****A****^p^*⁄*∂t*. Omitting the time-dependent scale factor, − *di_j_* ⁄*dt*, one has

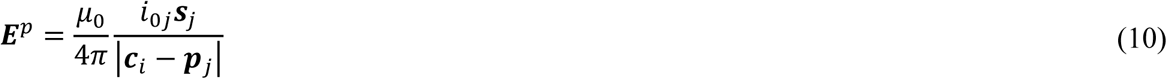

For every observation point, the primary electric field in Eq. (10) is computed via the FMM as a potential of a single layer repeated three times, i.e., separately for each component of the field. Examples of detailed TMS coil models are shown in Fig. 2; the coil geometry generator of the toolkit is described in Appendix A.

**Fig. 2.**
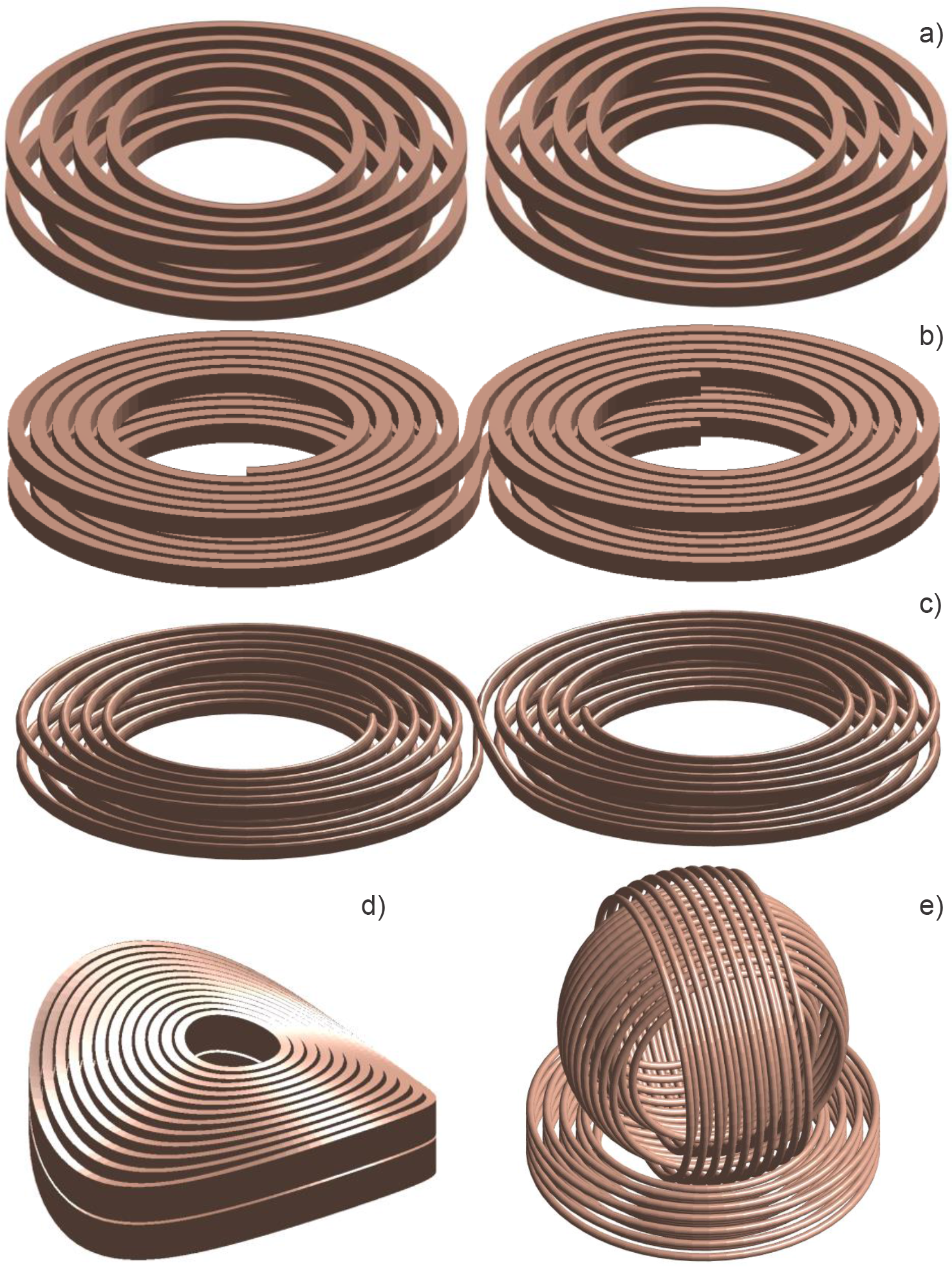
Some solid CAD models created using the MATLAB-based coil geometry generator. Fig. 2a is a simplified MRi-B91 TMS-MRI coil model (MagVenture, Denmark) with elliptical conductors of a rectangular cross-section used in this example; Fig. 2b is a simplified MagPro C-B60 coil model (MagVenture, Denmark); Fig. 2c is a generic double figure-eight spiral coil model with an elliptical cross-section and two bootstrapped interconnections; Fig. 2d is a simplified Cool-40 Rat small animal coil model (MagVenture, Denmark); Fig. 2e is a three-axis multichannel TMS coil array radiator (Navarro de Lara et al 2018).

### 2.9. Excitation

The excitation is the term on the right-hand side of Eq. (4) which depends on ***E****^p^*. It is evaluated first at mesh nodes via the FMM, and then an average value for every facet center is obtained. The initial guess of an iterative solution is proportional to the excitation term. Based on analytical solutions for the sphere and ellipsoid, a scalar weighting parameter for the initial guess may be chosen in the range from 1 to 10.

### 2.10. Iterative solution

In the iterative matrix-free solution, Eqs. (4) and (6) are added together and solved simultaneously. The weighting parameter for the conservation law normalized by the total surface area is chosen as 0.5. The generalized minimum residual method (GMRES) was found to converge better than the bi-conjugate gradients method and its variations. Several sparse near-field preconditioners have been constructed, but so far none has provided a significantly better convergence speed. The relative residual of 10^-10^ is achieved in approximately 60 iterations for a typical head model discussed further. However, such a large number of iterations is not necessary as shown below.

### 2.11. Discretization error and surface charge averaging

From the viewpoint of electromagnetic field theory (Van Bladel, 2007), any sharp edge in the surface mesh will lead to an infinite value of the surface charge density at that location, which indeed results in an infinite electric-field value when the mesh surrounding this edge is refined. Fortunately, this is an integrable singularity (Van Bladel, 2007). In order to compare models with different surface resolutions, we must therefore introduce averaging (integration) over a consistent and small surface area. For practical purposes, it is convenient to introduce equally weighted surface charge averaging (low pass filtering) over three or more immediate topological neighbors. After the solution is obtained, we substitute (for three topological neighbors)

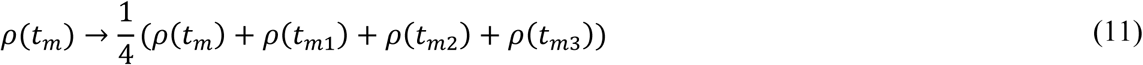

where triangles *t_mn_* share an edge with triangle *t_m_*. Eq. (11) has been implemented in the code. For a very fine mesh, it could be applied twice to expand the relative averaging domain.

### 2.12. Postprocessing

The normal electric field just inside/outside an interface is computed directly from the known surface charge density following Eqs. (5). The total electric field just inside/outside an interface follows Eq. (2) with all neighbor potential integrals computed analytically. The total electric field anywhere in the volume follows Eq. (1) with all neighbor potential integrals again computed analytically.

### 2.13. Available subject head models

Sixteen Human Connectome Project (HCP) head models (Van Essen et al 2012, 2019) with an initial isotropic voxel resolution of 0.7 mm are available with the software (Htet et al 2019b). These MRI data have been converted to surface models using the SimNIBS v2.1 pipeline; each model includes seven brain compartments (skin, skull, cerebrospinal fluid (CSF), gray matter (GM), white matter (WM), ventricles, cerebellum). Each model has been checked against the original NIfTI images, and mesh manifoldness has been strictly enforced and confirmed using ANSYS HFSS mesh checker. The default average cortical surface mesh edge length is 1.5 mm, the cortical nodal density is 0.55 nodes per mm^2^, and the total number of facets is 0.9 M.

Any other surface model may be used in *.stl or *.mat (MATLAB) format. A chief example is the MIDA model or its parts (Iacono et al 2015). Additionally, the fifty CAD models included in the Population Head Model Repository or PHM (Lee et al 2016, Lee et al 2018) may be used, which have been made available by the IT’IS Foundation, Switzerland via the web (IT’IS Foundation, 2016). Note that some inconsistencies were observed when overlapping these PHM models with the original NIfTI images.

### 2.14. Model remeshing and surface mesh registration

The MATLAB package also includes CM2 SurfRemesh^®^, a mesh generation program from Computing Objects, France, that enables the user to create coarser and/or finer surface representations while minimizing the surface deviation error from the master mesh. A NIfTI viewer available in the core MATLAB package facilitates surface mesh registration in any plane by overlapping surface mesh cross-sections on NIfTI images. We should note that most accurate registration the original MRI data header information should be used, and the provided viewer is predominantly to facilitate quick visualization in MATLAB.

### 2.15. Implementation and distribution

The complete BEM-FMM algorithm is implemented entirely in MATLAB^®^ 2019 for both Windows (runs as is) and Linux (may require machine-specific compilation of the FMM library). As mentioned above, the software includes the latest FMM library (Gimbutas et al. 2019) and is bundled together with remeshing and registration modules. The complete package is available to interested researchers via a Dropbox repository (Dropbox 2019).

## 3. Results

### 3.1. Necessary number of iterations for the iterative solution

A critical point for the both the speed and performance of the method is the number of GMRES iterations to be used in the iterative solution. This number, which determines the overall algorithm’s speed, was not quantified before.

In order to establish the representative estimate, six Connectome Project head models (subject numbers 110411, 117122, 120111, 122317, 122620, 124422, Van Essen et al 2012, 2019) and the corresponding surface models obtained via the SimNIBS v2.1 pipeline and described above were tested. The average cortical surface mesh edge length is 1.5 mm; the cortical nodal density is 0.55 nodes per mm^2^; the total number of facets is 0.9 M.

Further, the remeshing program described above was applied. As a result, a coarser set with an average cortical edge length of 1.9 mm and an average cortical nodal density of 0.32 nodes per mm^2^ was created; the total number of facets is 0.4 M. Likewise, a finer set with an average cortical edge length of 0.99 mm and a cortical nodal density of 1.2 nodes per mm^2^ was created; the total number of facets is 1.8 M. The models were augmented with the following material conductivities (at 3 kHz center frequency): scalp average – 0.333 S/m, skull – 0.0203 S/m, CSF –S/m, GM – 0.106 S/m, cerebellum – 0.126 S/m, WM – 0.065 S/m, ventricles – 2.0 S/m (Database of Tissue Properties. IT’IS Foundation 2019).

The widely used MRI compatible TMS coil MRi-B91 (MagVenture, Denmark) located above the motor hand area of the precentral gyrus (the hand knob area, Yousry et al 1997), was employed in these tests. A detailed coil model has been constructed and approximated by 26 thousand elementary current segments in Eq. (10). The coil was driven by a time-varying current of 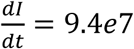 *Amperes/sec* The primary coil field was computed using Eq. (10).

In every case, the coil was positioned in order to follow three geometrical rules:

i. align the approximately identified hand knob area of the right precentral gyrus with the coil centerline;
ii. set the coil centerline approximately perpendicular to the skin surface, and position the coil 10±0.25 mm from the skin along this centerline;
iii. have the dominant field direction (the *x*-axis of SimNIBS coordinate system) approximately perpendicular to the gyral crown (and associated sulcal walls) of the precentral gyrus pattern at the target point.

These rules uniquely define the coil position and the rotation angles.

The most sensitive error parameter is the value of the absolute maximum field observed at the cortical interfaces and the exact position of this local maximum. Figs. 3 a,b,c show the error in the maximum value of the total field just inside the pial cortical surface or the gray matter shell (red) and just inside the inner cortical surface or the white matter shell (blue) as a function of iteration number versus the most accurate solution with 100 iterations for the three different model resolutions described above. Figs. 3 d,e,f give the error in the position of this maximum field in millimeters as a function of iteration number versus the most precise solution for the same three cases. The vertical line in every plot corresponds to the 15^th^ iteration. Results are only given for subject #110411 of the Connectome Project Database, but nearly identical results have been observed for the five other subjects considered. Based on these results we conclude that 14-15 iterations are sufficient to obtain a maximum-field error below 1% and a maximum-field position error below 0.25 mm.

Table 1 gives the error in the maximum value of the total field just inside the inner cortical surface and the error in the position of this maximum field in millimeters versus the most accurate solution available (with 1.2 nodes/mm^2^, 1.8 M facets, 100 iterations) for the two lower model resolutions. Results are given for all six subjects listed above. This surface-charge averaging area in Eq. (11) has been defined as 3 mm^2^. One can see that both errors are relatively small, even for the coarse model with 0.4 M facets.

**Table 1.** Error in the maximum value of the total field just inside the inner cortical surface (the white matter shell) and the error in the position of this maximum field in millimeters versus the more precise solution (with 1.2 nodes/mm^2^, 1.8 M facets) for two lower model resolutions. Results are given for six subjects of the Connectome Project Database.

| **Connectome subject** | **Max field error for 1.5 mm edge length (0.55 nodes/mm<sup>2</sup>, 0.9 M facets)** | **Error in the position of the max field (0.55 nodes/mm<sup>2</sup>, 0.9 M facets)** | **Max field error for 1.9 mm edge length (0.32 nodes/mm<sup>2</sup>, 0.4 M facets)** | **Error in the position of the max field (0.32 nodes/mm<sup>2</sup>, 0.4 M facets)** |
| --- | --- | --- | --- | --- |
| 110411 | 0.6% | 0.2 mm | 3.8% | 0.9 mm |
| 117122 | 1.2% | 0.1 mm | 1.2% | 1.7 mm |
| 120111 | 0.9% | 0.2 mm | 0.2% | 1.1 mm |
| 122317 | 0.3% | 0.7 mm | 2.3% | 1.5 mm |
| 122620 | 0.5% | 1.1 mm | 2.1% | 0.9 mm |
| 124422 | 2.2% | 0.6 mm | 3.5% | 0.8 mm |
| **Average** | **0.9 %** | **0.5 mm** | **2.2%** | **1.2 mm** |

### 3.2. Overall method speed

Based on results shown in Fig. 3 and quantified in Table 1, we estimate the speed of the method sufficient for an accurate solution pertinent to different model resolutions in Table 2. A significant speed improvement by a factor of approximately 3 as compared with previous results (Makarov et al 2018, Htet et al 2019a) is achieved by using an improved FMM algorithm, by employing the proper number of iterations, by lowering the intrinsic FMM precision to an acceptable level without compromising the overall method accuracy, and by the explicit inclusion of the charge conservation law into the iterative solution. The last step provides an excellent convergence of the iterative solution in all cases considered detailed in Appendix A.

**Fig. 3.**
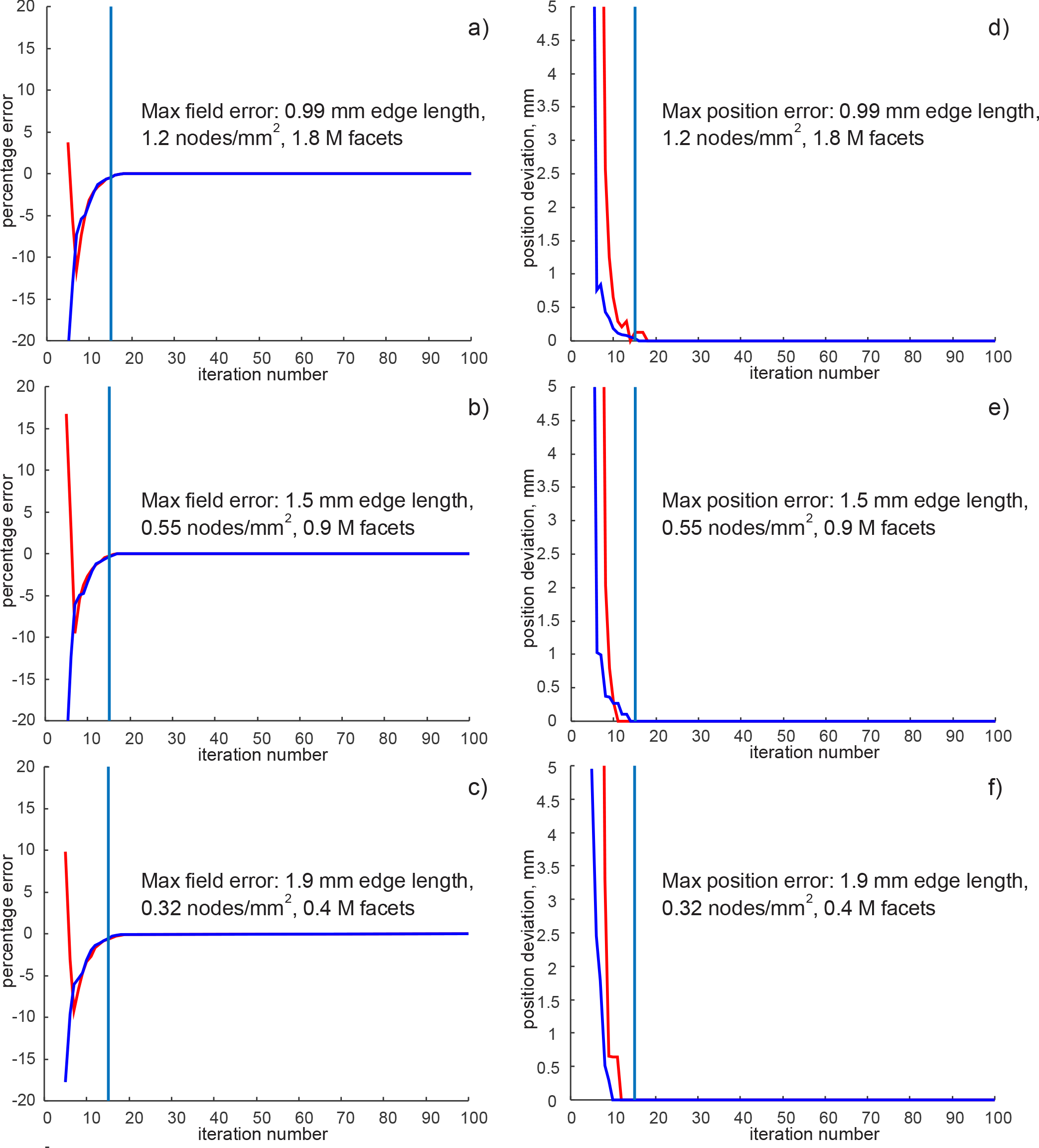
a,b,c) – Error in the maximum value of the total field just inside the gray matter shell (red) and white matter shell (blue) as a function of iteration number versus the most accurate solution with 100 iterations for three different model resolutions. d,e,f) – Error in the position of this maximum field in millimeters as a function of iteration number versus the most precise solution for three different model resolutions. The vertical line in each plot corresponds to the 15^th^ iteration. Results are given for subject #110411 of the Connectome Project Database. Very similar results were observed for the five other subjects considered.

**Table 2.** BEM-FMM run times for different model resolutions obtained with 14 iterations. These data were compiled using an Intel Xeon E5-2698 v4 CPU (2.10 GHz) server, 256 GB RAM, MATLAB 2019a.

| **Model resolution** | **Preprocessing time (once per model)** | **Solution time for 14 iterations (cf. Fig. 1)** | **Post processing time for normal surface cortical fields and field discontinuities\*)** |
| --- | --- | --- | --- |
| 1.2 nodes/mm<sup>2</sup>, 1.8 M facets | 110 sec | 74 sec | 0 sec |
| 0.55 nodes/mm<sup>2</sup>, 0.9 M facets | 70 sec | 34 sec | 0 sec |
| 0.32 nodes/mm<sup>2</sup>, 0.4 M facets | 50 sec | 18 sec | 0 sec |

\*) Post-processing time for the total/tangential interfacial fields should also approach zero once the last GMRES iterations are made available in the workspace.

### 3.3. Modeling example: computing total, tangential, and normal fields at the interfaces

The developed algorithm is applied to accurately compute normal, tangential, and total electric fields anywhere in the cerebral cortex for a specific subject and a specific coil orientation. In this example, particular attention is paid to modeling the field in the vicinity of the folded white-gray matter interface (the inner cortical surface).

Electric fields in the brain were simulated using the MRi-B91 coil model, located above the motor hand area of the right hemisphere in six distinct subject models of the Connectome Project described above in section 3.1. Only the finer resolution models with an average cortical edge length of 0.99 mm, a cortical nodal density of 1.2 nodes per mm^2^, and a total number of facets of 1.8 M have been used for simulations. We employed straightforward geometrical coil positioning as described in section 3.1 above.

Fig. 4a shows the computed magnitude of the total surface electric field for subject #110411 just inside the pial cortical surface (the gray matter shell). This is the typical non-focal gray-matter field distribution observed in many relevant studies. The field distribution includes a number of sparse local maxima, one of which is located close to the coil centerline. The absolute field maximum for the plot is 151.5 V/m. In general, the field just inside the gray matter shell corresponds to cortical layer I, which is likely of little interest for TMS activation (Thielscher 2019).

A different situation occurs when we evaluate the total field magnitude just inside the inner cortical surface or the white matter shell, as shown in Fig. 4b. Here, the absolute field maximum is somewhat lower and computed to be 138.6 V/m. However, the field has become quite focal; the maximum field is concentrated in a well-defined domain marked by an arrow in Fig. 4b.

**Fig. 4.**
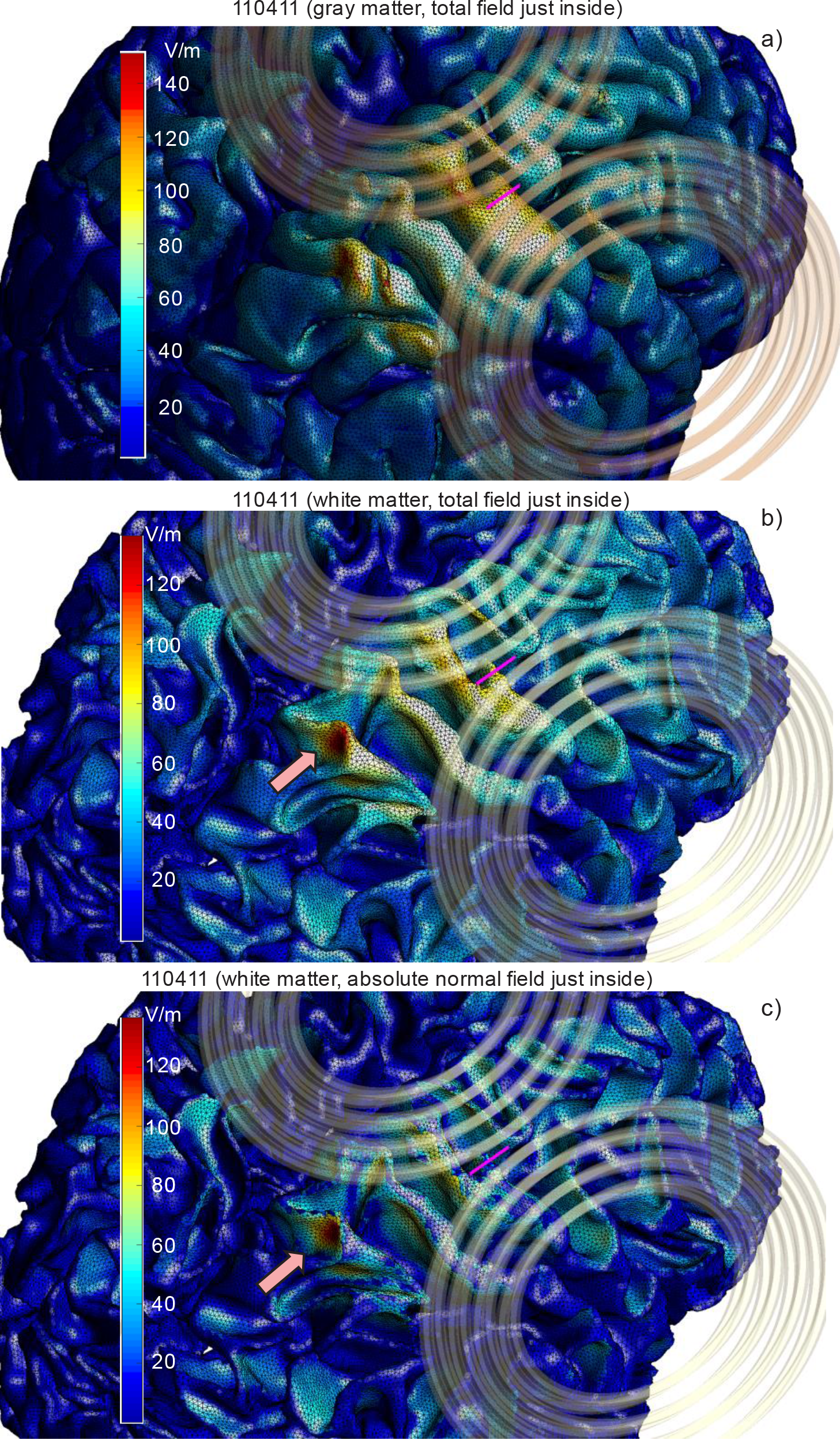
a) – Computed total surface electric field (magnitude) distribution just inside the pial surface (gray matter shell) for subject #110411. The absolute field maximum for the plot is 151.5 V/m. b) – The same result just inside the white matter shell; the absolute field maximum is 138.6 V/m. c) The normal-field magnitude just inside the white matter shell; the field maximum is 134.2 V/m.

Remarkably, this focal domain resides in the area of the superior parietal lobule, just behind the postcentral sulcus and rather far away from the targeted hand knob area of the precentral gyrus. The distance from the coil centerline intersection with the white matter shell to the center of the depicted hot spot is equal to 32 mm. The distance from the coil centerline intersection with the GM shell is even longer. This observation is consistent with the previously established fact that the apparent TMS motor map may extend due to remote hotspot activation (Reijonen et al 2019).

Which field component generates this local maximum? To answer this question, Fig. 4c plots the absolute value of the normal field just inside the white matter shell. In this instance, the focal area is even more pronounced with an absolute field maximum of 134.2 V/m. Comparing this number with the previous value of 138.6 V/m, we conclude that the normal field component is primarily responsible for this maximum.

### 3.4. Modeling example: computing normal fields and their discontinuities at the white-gray matter interface

The normal field to the inner cortical surface (the white-gray matter interface) is the field parallel to the long, either straight or slightly bent pyramidal axons of the fast-conducting pyramidal tract neurons passing through this interface. The normal field discontinuity (or, rather, a very rapid field variation) across the white-gray matter interface creates perhaps the strongest gradient of the component of the electric field along the axon, excluding the effect of the axonal bending (Miranda et al 2007). It has been suggested that such a gradient may cause stimulation (Miranda et al 2007, Salvador et al 2011). The stimulation of pyramidal axons of the fast-conducting pyramidal tract neurons results in D (direct) TMS wave generation (Salvador et al 2011, Lazzaro and Ziemann 2013), which can be measured experimentally (Lazzaro and Ziemann 2013).

Fig. 5 shows the white-gray matter interfaces (the white matter shells) for six Connectome subjects considered. Small blue spheres are drawn at the center of every white matter facet where the absolute normal field value just inside the surface is in the range of 80-100% of the maximum normal-field value for the same surface. Identical “hot spots” would also be observed for the normal component just outside the interface and for the normal field discontinuity across the interface, according to Eq. (5).

**Fig. 5.**
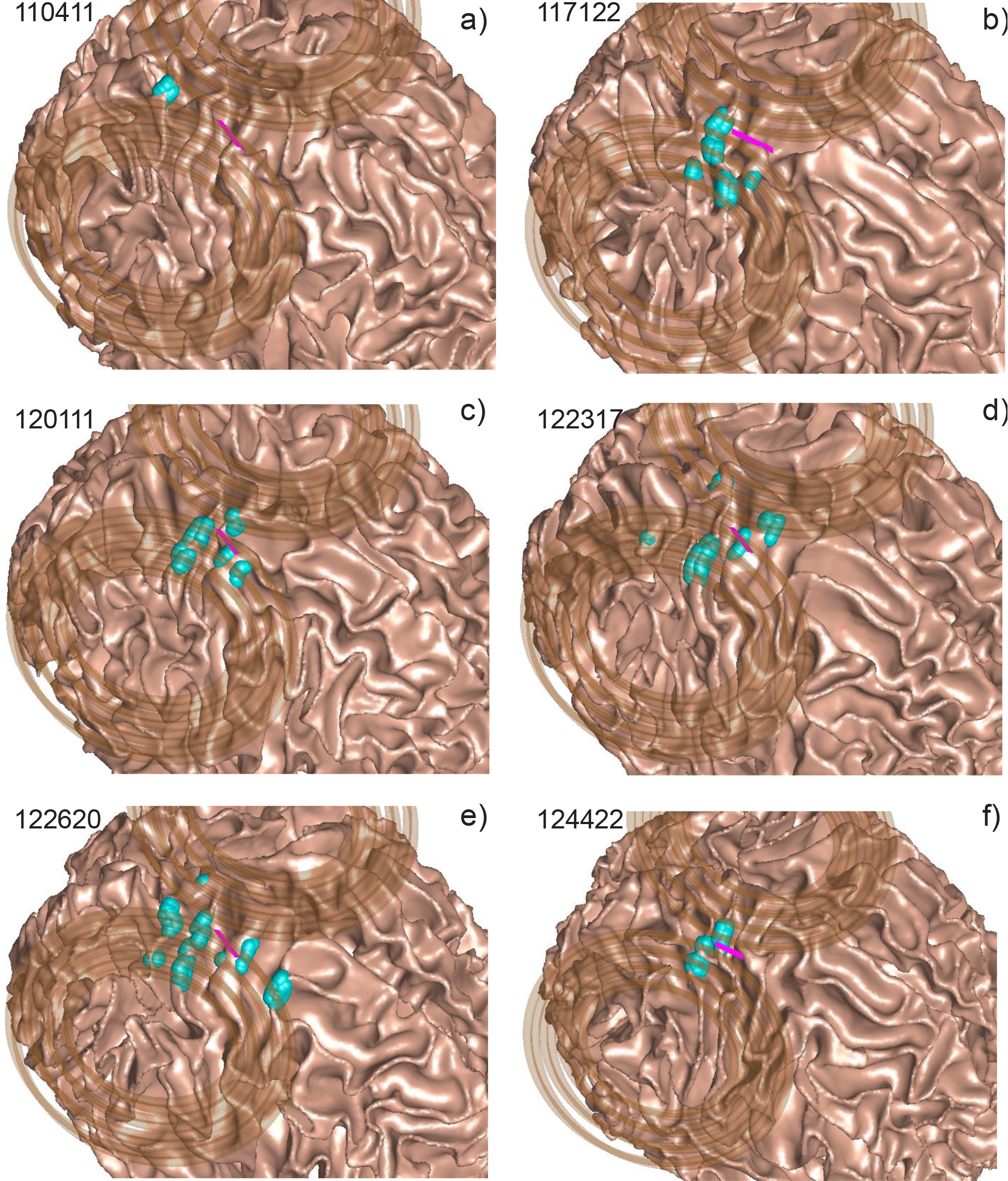
Distribution of E-field “hot spots” for the normal-field component at the inner cortical surfaces using geometrical coil positioning. a)– f) – Inner cortical surfaces (white-gray matter interfaces) for six Connectome subjects with small blue balls drawn at the center of every white matter facet where the absolute normal-field value just inside the surface is in the range of 80-100% of the maximum normal-field value for the same surface. In every case, the MagVenture MRiB91 coil is targeting the motor hand area of the right hemisphere via geometrical positioning. The total WM areas with the normal-field values within 80-100% of the maximum normal field are 43, 119, 120, 170, 197, and 82 mm^2^; the standard deviation of all high-field values from the maximum-field position is 2.4, 9.0, 8.5, 12, 17, and 6.6 mm for Connectome subjects 110411, 117122, 120111, 122317, 122620, and 124422 respectively.

In every case, the TMS coil is targeting the motor hand area of the right hemisphere. The total WM areas with the normal-field values within 80-100% of the maximum normal-field value are 43, 119, 120, 170, 197, and 82 mm^2^; the standard deviations of all high-field values from the maximum-field position are 2.4, 9.0, 8.5, 12, 17, and 6.6 mm for the six Connectome subjects 110411, 117122, 120111, 122317, 122620, and 124422 respectively.

One can see that, in the majority of cases in Fig. 5, the TMS response with regard to the normal inner cortical field becomes sparse and often significantly deviates from the coil centerline. The intersubject variations are also strong.

In this example, the most remarkable result has been observed for subject #110411 (Fig. 5a). The total WM area covering 80-100% of the maximum field in Fig. 5a is compact; its size is only 43 mm^2^ (approximately 6.6 mm × 6.6 mm). According to Eq. (5), the same focal area is observed for both the normal field just outside the inner cortical surface and for the normal field discontinuity across the inner cortical surface. Note again that the maximum values of the field just inside and outside the white matter shell are 134.2 V/m and 82.3 V/m, respectively, demonstrating significantly higher field values just inside the inner cortical surface and a large field discontinuity. This result directly follows from Eq. (5) when the corresponding conductivity values are substituted.

A separate study has been performed to test if the result for subject #110411 is stable with respect to perturbations in the coil position. In order to accomplish this, the algorithm was straightforwardly modified to run in a parametric loop with no graphical output. It was found that, if the variations in the coil rotation angles and in three coil coordinates do not each exceed 3%, both the focal position and the focal field value remain stable.

### 3.5. Modeling example: computing high-resolution volumetric field distributions

To demonstrate the BEM-FMM algorithm’s field resolution capability, we consider the region in the vicinity of the E-field maximum in Fig. 5a, where the electric field likely changes very rapidly. The corresponding results for three principal planes passing through the field maximum position in Fig. 5a are given in Figs. 6, 7, and 8. In Figs. 6a, 7a, and 8a, the surface mesh has been overlaid on top of the relevant NIfTI slice to demonstrate mesh registration with the original imaging data. The red dots again signify the centers of intersected white matter facets where the absolute normal-field value is in the range of 80-100% of the maximum normal-field value. Field localization in these planes is very good.

**Fig. 6.**
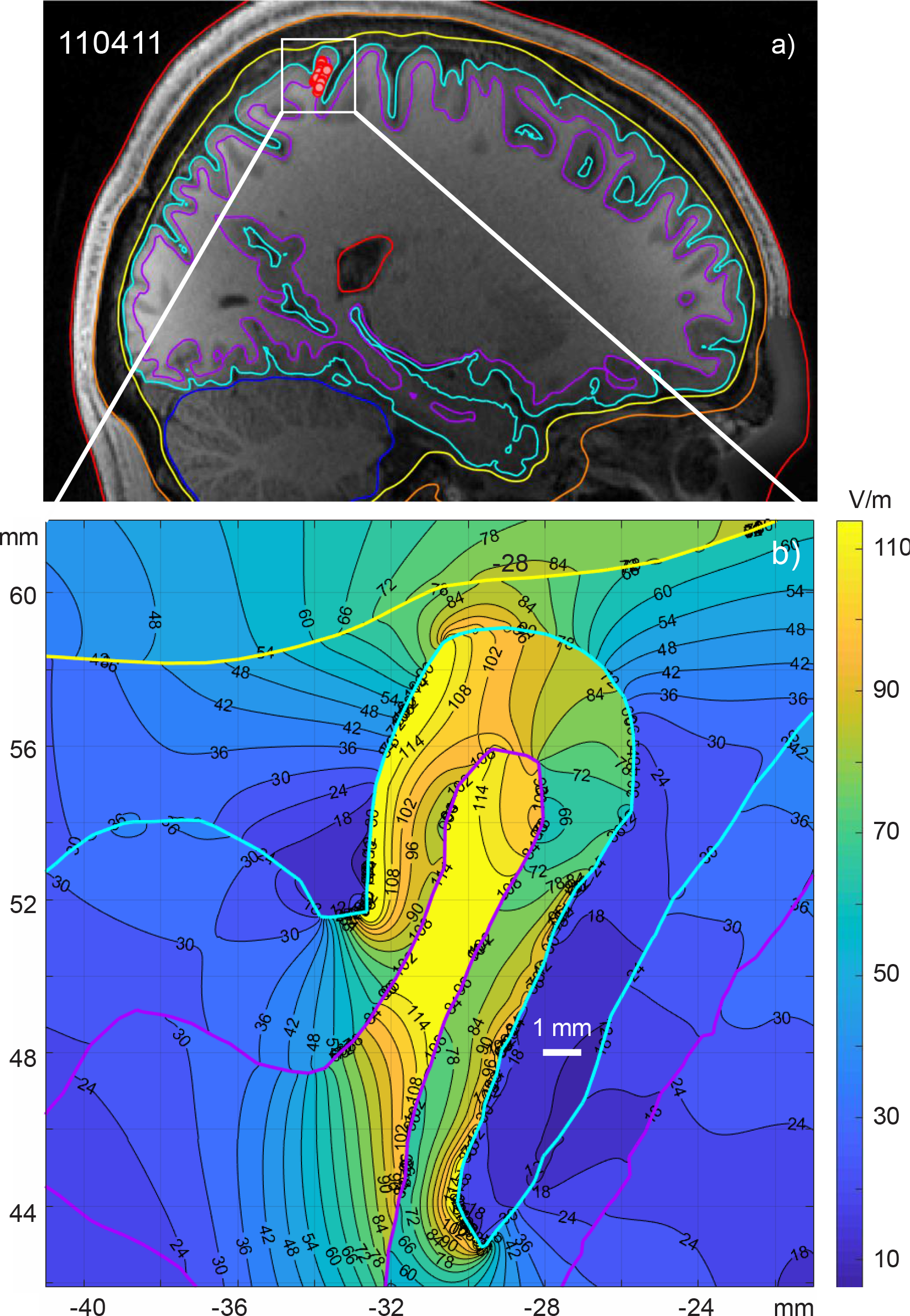
a) – Sagittal plane passing through the location of the maximum field at the white-gray matter interface and superimposed onto the corresponding NIfTI slices when using the MagVenture MRiB91 coil located above the hand knob area of the right precentral gyrus of subject #110411. The red dots indicate the centers of intersected white matter facets where the absolute field value is in the range of 80-100% of the maximum field value. b) – volumetric total field (magnitude) distribution within the small white rectangle in Fig. 6a. Blue color – CSF-gray matter interface, purple color – white-gray matter interface, yellow color – skull-CSF interface.

**Fig. 7.**
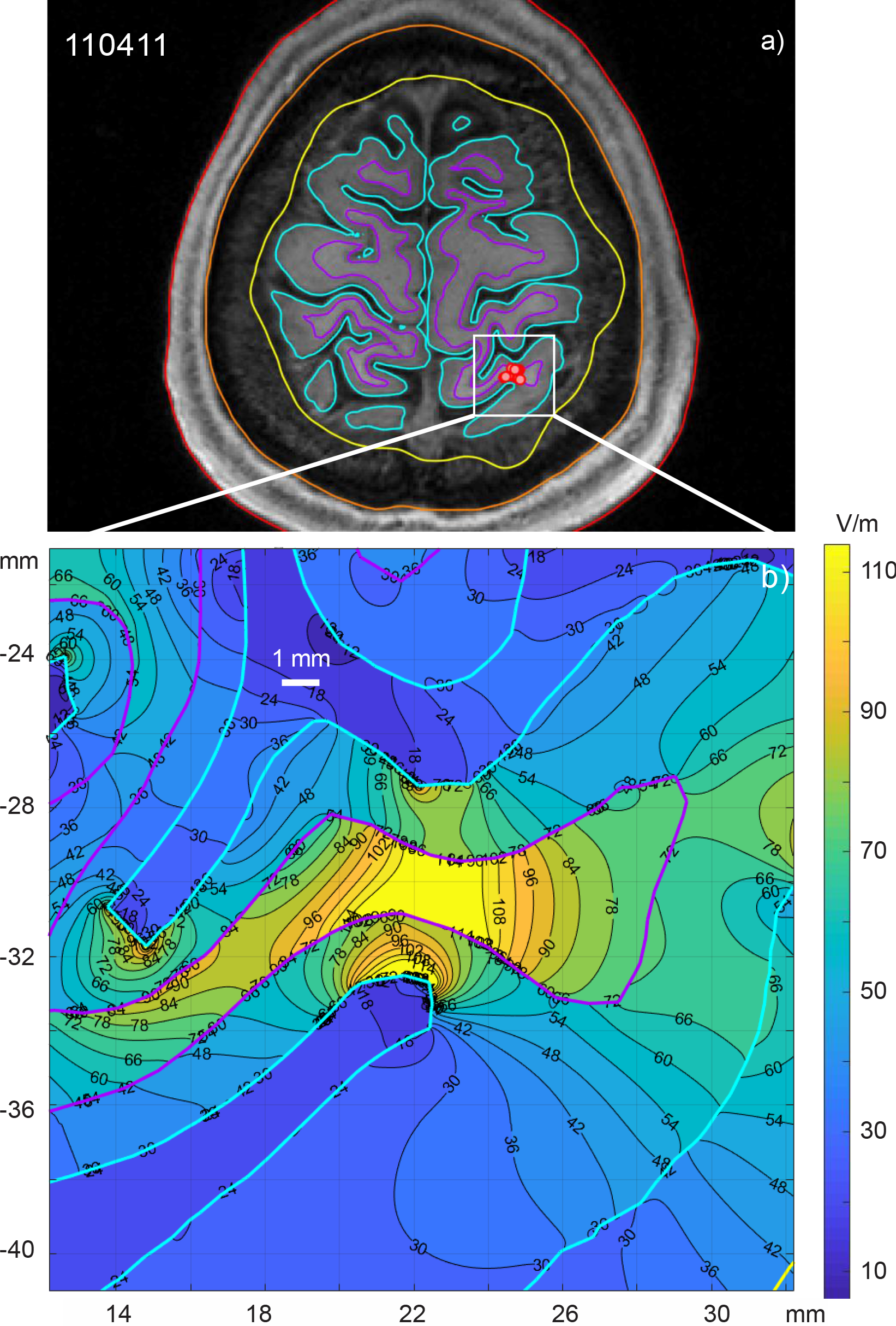
a) – Transverse plane passing through the location of the maximum field at the white-gray matter interface and superimposed onto the corresponding NIfTI slices for the MagVenture MRiB91 coil located above the hand knob area of the right precentral gyrus of subject #110411. The red dots depict the centers of intersected white matter facets where the absolute field value is in the range of 80-100% of the maximum field value. b) – volumetric total field (magnitude) distribution within the small white rectangle in Fig. 7a. The same notations from Fig. 6 are used.

**Fig. 8.**
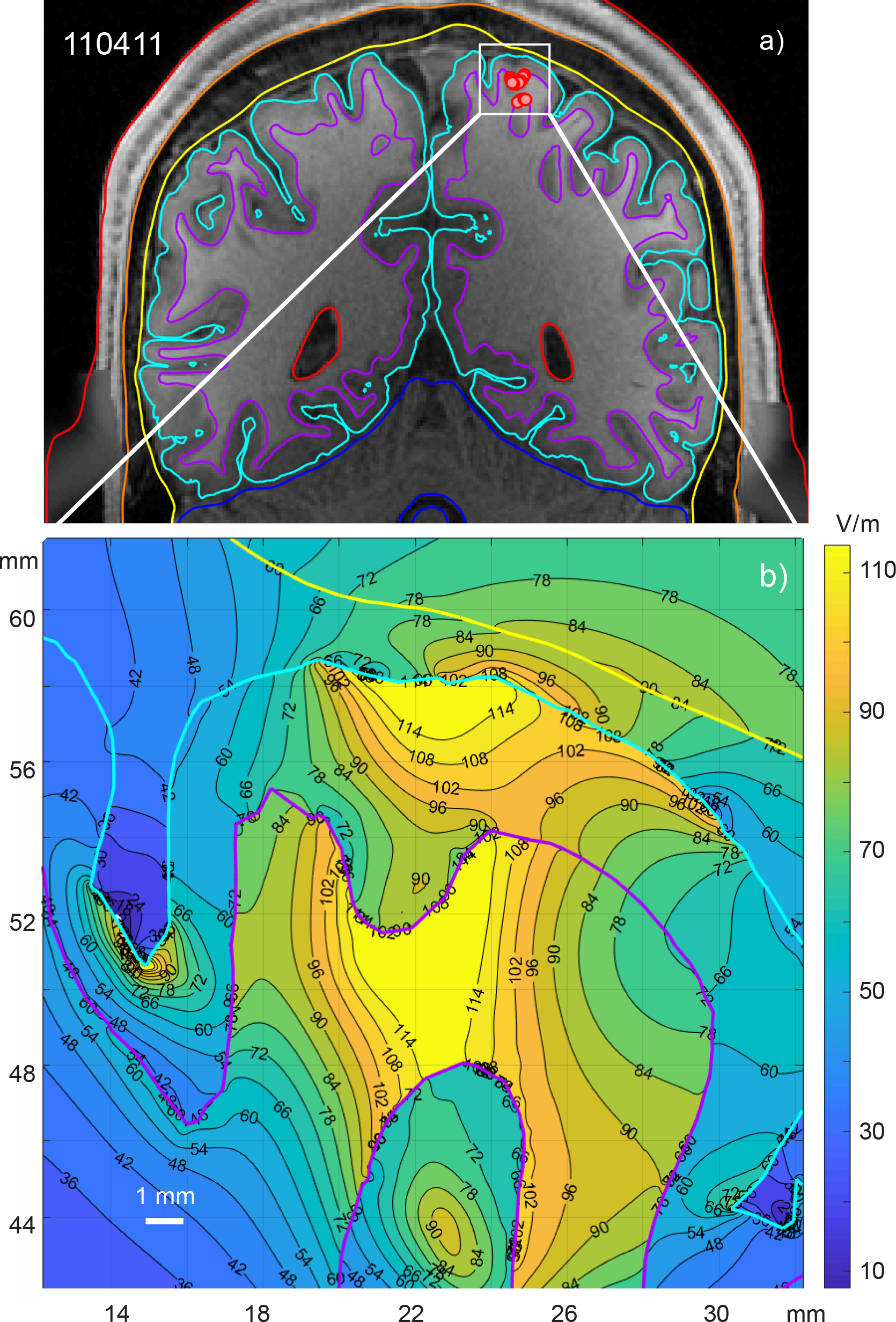
a) – Coronal plane passing through the location of the maximum field at the white-gray matter interface and superimposed onto the corresponding NIfTI slices for the MagVenture MRiB91 coil located above the hand knob area of the right precentral gyrus of subject #110411. The red dots show the centers of intersected white matter facets where the absolute field value is in the range of 80-100% of the maximum field value. b) – volumetric total field (magnitude) distribution within the small white rectangle in Fig. 8a. The same notations from Fig. 6 are used.

The volumetric field distribution in Figs. 6b, 7b, and 8b was obtained in the areas labeled by white rectangles in Fig. 6a, 7a, and 8a. Each area is 20×20 mm^2^ and is centered exactly at the location of the absolute field maximum. Each area contains 500×500 observation points resulting in a field resolution of 40 µm. The volumetric field distribution is given for the magnitude of the *total* electric field. Specific colors designate interfaces: blue corresponds to the pial cortical surface, purple defines the inner cortical surface, and yellow defines the skull-CSF interface.

We see in Figs. 6b, 7b, and 8b that the local volumetric field inside the narrow white matter gyrus changes very rapidly, at the submillimeter scales. The same is valid for the surrounding gray matter. The volumetric divergence of the electric field or its one-dimensional counterpart, the activating function *dE*/*dz*, easily reaches 1-10 V/mm^2^ in both the white matter gyral crown/lip and in the white matter sulcal walls.

## 4. Discussion

### 4.1. Comparison with analytical solutions and with other software packages/numerical methods

Three separate computational studies (Makarov et al 2018, Htet et al 2019a, Gomez et al 2019) have been performed to compare the method’s performance in application to TMS problems. For the canonic multisphere geometry and an external magnetic-dipole excitation where the analytical solution is available, the BEM-FMM algorithm was tested against a fast open-source getDP solver running within the SimNIBS 2.1.1 environment (Htet et al 2019a). It was observed that the BEM-FMM algorithm gives a smaller solution error for all mesh resolutions and runs significantly faster for high-resolution meshes when the number of triangular facets exceeds approximately 0.25 M (Htet et al 2019a). The algorithm was further tested for 10 realistic head models of Population Head Model Repository (Lee et al 2016, Lee et al 2018) excited by a realistic coil geometry. The algorithm’s performance was compared against a high-end commercial FEM software package ANSYS Maxwell 3D with adaptive mesh refinement (Makarov et al 2018). Excellent agreement was observed for electric field distribution across different intracranial compartments, and the BEM-FMM algorithm achieved a speed improvement of three orders of magnitude over the commercial FEM software. (Htet et al 2019a).

A detailed and rigorous comparison study was recently performed independently by another group (Gomez et al 2019). For MRI-derived head models, the method of the present study – the 0th order BEM-FMM – was determined to be the most accurate method that could be run with available computational resources. Other methods (from least to most accurate, Gomez et al 2019): FDM or finite difference method, 1st order FEM, SPR (superconvergent patch recovery)-FEM, 2nd order FEM, 1st order BEM, and 3rd order FEM were benchmarked against the present method. It was concluded that, whereas at present the 1st order FEM is most commonly used, the 0th order BEM-FMM appears to be the judicious strategy for achieving negligible numerical error relative to modeling error, while maintaining tractable levels of computation.

### 4.2. Method speed and model size

The algorithm runs best on multicore workstations/servers and multicore PCs due to the inherent parallelization of MATLAB and the FMM package. The number of cores seems to be more important than the clock speed. According to Table 2, the numerical TMS solution for the head segmentation with 0.55 nodes/mm^2^ and 0.9 M facets in total executes with the improved BEM-FMM in approximately 34 sec using a 2.1 GHz multicore server. This is threefold improvement compared to the initial formulation (Makarov et al 2018 or Htet et al 2019a). The numerical TMS solution with 1.8 M facets in total from Table 2 executes in approximately 74 sec, i.e. scales nearly linearly with the number of facets.

A surface model with 70 M facets has been considered and computed with the present toolkit. For the same 2.1 GHz multicore server listed in Table 2, the corresponding execution time reaches approximately two hours.

However, for a laptop computer, the BEM-FMM algorithm is expected to run significantly slower than the FEM pipeline of SimNIBS 3.0 (Saturnino et al 2019c) when a comparable resolution model is used. The BEM-FMM algorithm is also, in the present version, unable to handle white matter anisotropy.

Also, the BEM-FMM approach requires relatively large preprocessing times for an individualized head model in order to compute and store necessary neighbor electrostatic potential integrals on triangular facets. Although this preprocessing step is required only once, the time required is on the order of one minute or longer.

### 4.3. Volumetric field resolution

In contrast to FEM, the BEM-FMM resolution in the cortex is not limited by the volumetric tetrahedral mesh size; it may reach a micron scale if necessary. The key difference is that the solution is completely determined by the incident field and the induced charge distribution on the conductivity boundary surfaces, and the E-fields at arbitrary points of the 3D space can be subsequently evaluated. In the present study, we have demonstrated the ability to accurately compute TMS fields within the cortex at submillimeter scales with a field resolution of 40 µm as well as close to and across the cortical interfaces, in particular across the white-gray matter interface.

The method accuracy is only limited by the surface segmentation itself and not by the volumetric mesh size. A meaningful solution is obtained at any distance from the interface, including a distance approaching zero. For example, for two interfaces separated by 2 mm, BEM-FMM is expected to generate more accurate results than the first-order FEM of SimNIBS at the distances of 0.5 mm or less from the either interface unless many tetrahedra across the 2 mm gap or higher-order FEM basis functions are used. Indeed, the segmentation accuracy itself provides a limit on the overall modeling accuracy. However, detailed segmented models with isotropic resolution of 0.5 mm are already available (Iacono et al 2015).

Furthermore, it is seen in Figs. 6, 7, 8 that the field in the cortex may change very rapidly and at submillimeter scales. The accurate high-resolution field modeling may therefore be important for subsequent multiscale modeling pertinent to evaluating the neuron activating function (Wang et al 2018, Aberra et al 2018, Aberra et al 2019).

### 4.4. Interfacial field resolution

The electric field is discontinuous at the interfaces of brain compartments with different conductivities, due to surface charge accumulation and the abrupt jump of the normal field component across a single monopolar layer of charges. In the present study, we distinguish the normal field just inside the interface, the normal field just outside, and the normal field discontinuity across the interface. The BEM-FMM approach (and any BEM method) accurately accounts for the interface field discontinuities. Some FEM solvers (e.g., ANSYS Maxwell, see Htet et al 2019a) also resolve these discontinuities precisely whereas others (for example, SimNIBS) may perform spatial smoothing instead.

It should be noted that the infinitely-thin white-gray matter interface is a physical modeling assumption of the underlying biological tissue structure, thus raising the question: is the mathematical field discontinuity relevant? For example, the myelination that is important for the electrical insulation of the axons, starts to increase already in cortical layer IV and below in the rat neocortex (Srinivasan et al 2012). From a formal point of view, the macroscopic field discontinuity leads to infinite values of the activating function, proportional to *dE*/*dz*, for pyramidal axons passing through the white-gray matter interface with the normal coordinate *z*. The BEM-FMM approach correctly computes the finite difference, *dE*, following Eq. (5), but *dz*, the “effective” thickness of the white-gray matter interface, remains unknown for this and other macroscopic methods where it is set to zero.

Fortunately, a solution of the cable equation with the abrupt field discontinuity does exist; it predicts the finite values of the transmembrane potential proportional to *dE*(Miranda et al 2007). This solution might perhaps contain more meaningful information than *dE*/*dz* found on the base of spatial smoothing, which depends on the non-biological numerical parameter: the size and the local density of a particular FEM computational tetrahedral mesh. For example, the field discontinuity of 52 V/m observed in Fig. 5a induces a transmembrane potential of 52 mV for a straight pyramidal axon perpendicular to the white-gray matter interface if the membrane space constant of Miranda et al 2007 is used.

## 5. Conclusions

In this study, we have described the improved BEM-FMM numerical algorithm for TMS modeling. Compared with previous results, the BEM-FMM algorithm has been improved in several novel ways. First, we have established a fast, non-saturated solution convergence by incorporating an explicit global charge conservation law. Second, we utilized a simple analytical approach for obtaining electric fields (and electric field discontinuities) normal to the cortical surface (or any other interfaces) at no extra computational cost and without any postprocessing or smoothing. Third, we established a minimum sufficient number of iterations for obtaining an accurate solution. Finally, we have incorporated a fully general treatment of the boundary interface geometries, allowing non-nested surface models to be used.

The improvements have increased the method speed by a factor of approximately 3, while maintaining the same accuracy, and have provided fast non-saturated convergence to arbitrarily small values of the relative residual (Appendix A). The numerical TMS solution for the head segmentation with 0.55 nodes/mm^2^ and 0.9 M facets in total now executes in approximately 34 sec using a 2.1 GHz multicore server.

The algorithm is based on the new general-purpose FMM kernel developed by the group of L. Greengard (Gimbutas et al 2019). The algorithm, coupled with tools that support surface remeshing and registration with corresponding NIfTI data, is implemented entirely in MATLAB and employs a few necessary toolboxes. It runs best on multicore machines due to the inherent parallelization of its platform. The number of cores seems to be more important than the clock speed. The complete computational code for this study, along with supporting documentation (Appendix A), is available online via a Dropbox repository (Dropbox, 2019).

In the present study, we have demonstrated the ability to accurately compute TMS fields within the cortex at submillimeter scales as well as close to and across the cortical surfaces, in particular across the white-gray matter interface. The method accuracy is only limited by the surface meshing itself and not by the nominal volumetric resolution of the MRI data. A meaningful solution is obtained at any distance from the conductivity boundary, including a distance approaching zero. It was found that it is possible to scan through electric fields normal to interfaces in real time and without postprocessing. The same functionality is true for the tangential and total interface fields once an intermediate integral in Eq. (2) is computed or acquired from the last step of the iterative solution.

The computational method developed may be useful for navigated TMS (Schmidt et al 2015, Fang et al 2019, Reijonen et al 2019) and robotic TMS systems (Goetz et al 2019) operating with the guidance of available high-resolution MRI imaging and providing accurate and stable coil position and orientation.

The mention of commercial products, their sources, or their use in connection with material reported herein is not to be construed as either an actual or implied endorsement of such products by the Department of Health and Human Services.

## Acknowledgements

The authors wish to thank Dr. Leslie Greengard and Dr. Manas Rachh of the Flatiron Inst., Ctr. for Computational Mathematics, NYC, USA for their continuous support of the fast multipole method implementation and the corresponding improvements. The authors are thankful to Dr. Maurie Klee for the useful discussion on the role of surface charge density in better understanding the cortical electric fields. The authors also would to thank Dr. Axel Thielscher and Mr. Guilherme Saturnino of Danish Research Center for Magnetic Resonance, Copenhagen University Hospital Hvidovre, Denmark for numerous useful discussions. This work has been partially supported by the National Institutes of Health (NIH) grants R00EB015445, NINDS R44NS090894, and R01MH111829.

